# Repurposing beta3-adrenergic receptor agonists for Alzheimer’s disease: Beneficial effects on recognition memory and amyloid pathology in a mouse model

**DOI:** 10.1101/2020.05.25.114454

**Authors:** Marine Tournissac, Tra-My Vu, Nika Vrabic, Clara Hozer, Cyntia Tremblay, Koralie Mélançon, Emmanuel Planel, Fabien Pifferi, Frédéric Calon

**Affiliations:** Faculté de pharmacie, Université Laval, Québec, Qc, Canada; Axe Neurosciences, Centre de recherche du CHU de Québec-Université Laval, Québec, Qc, Canada; UMR 7179 Centre National de la Recherche Scientifique, Muséum National d’Histoire Naturelle, Brunoy, France; Département de psychiatrie et neurosciences, Faculté de médecine, Université Laval, Québec, Qc, Canada

**Author notes:** Corresponding Author: Frederic Calon, Centre de recherche du CHU de Québec-Université Laval (Pavillon CHUL), Room T-2-67, 2705 boul. Laurier, Québec (Québec) G1V 4G2.

**Keywords:** Alzheimer’s disease, β3-adrenergic receptors, drug repurposing, 3xTg-AD mice, brown adipose tissue, thermogenesis

## Abstract

Old age, the most important risk factor for Alzheimer’s disease (AD), is associated with thermoregulatory deficits. Brown adipose tissue (BAT) is the main thermogenic driver in mammals and its stimulation, through β3-adrenergic receptor (β3AR) agonists or cold acclimation, counteracts metabolic deficits in rodents and humans. Studies in animal models show that AD neuropathology leads to thermoregulatory deficits and cold-induced tau hyperphosphorylation is prevented by BAT stimulation through cold acclimation. Since metabolic disorders and AD share strong pathogenic links, we hypothesized that BAT stimulation through a β3AR agonist could exert benefits in AD as well.

Here, we show that β3AR agonist administration (CL-316,243, 1 mg/kg/day i.p., from 15 to 16 months of age) decreased body weight and improved peripheral glucose metabolism and BAT thermogenesis in both non-transgenic and 3xTg-AD mice. One-month treatment with a β3AR agonist increased recognition index by 19% in 16-month-old 3xTg-AD mice compared to pre-treatment (14-month-old). Locomotion, anxiety and tau pathology were not modified. Finally, insoluble Aβ42/Aβ40 ratio was decreased by 27% in the hippocampus of CL-316,243-injected 3xTg-AD mice.

Overall, our results indicate that β3AR stimulation reverses memory deficits and shifts downward the insoluble Aβ42/Aβ40 ratio in 16-month-old 3xTg-AD mice. As β3AR agonists are being clinically developed for metabolic disorders, repurposing them in AD could be a valuable therapeutic strategy.

## INTRODUCTION

Old age is the main risk factor of Alzheimer’s disease (AD), a neurodegenerative disorder clinically expressed by memory deficits and cognitive dysfunction (Querfurth and LaFerla, 2010; Scheltens et al., 2016). The prevalence of AD is growing fast along with the aging population (Winblad et al., 2016). Yet, the exact pathogenic causes of the sporadic form of the disease are unknown. Despite decades of intense research and clinical trials, there is still no curative treatment for AD. Since AD is a complex and multifactorial disease, with frequent age-related comorbidities, multi-target agents might be advantageous over a single-bullet approach. The undeniable impact of old age on AD incidence indicates that aging triggers etiopathological factors of AD; identifying these key factors could provide invaluable clues to the development of novel therapeutic treatments.

Deficits in thermoregulation are among the documented consequences of old age. Although few studies investigated thermoregulation in AD individuals, it is well known that thermoregulatory defects appear in the elderly, the population primarily affected by AD (Degroot and Kenney, 2007; Gomolin et al., 2005; Grassi et al., 2003; Hebert et al., 2013). Mounting evidence now supports the hypothesis that thermoregulation deficits contribute to the development of AD pathology. Spontaneous thermoregulation deficits occurs in mouse models of AD neuropathology, including the triple transgenic (3xTg-AD) mice (Huitrón-Reséndiz et al., 2002; Knight et al., 2013; Sterniczuk et al., 2010; Vandal et al., 2016). Studies in mouse and hibernators repeatedly showed that decreased body temperature leads to increased tau phosphorylation (Arendt et al., 2003; Planel et al., 2007; Stieler et al., 2011). Accordingly, manipulation of external temperature leads to strong modulation of AD neuropathology in mice: cold exposure increases both tau phosphorylation and β-amyloid (Aβ) pathology and decreases synaptic proteins, while higher temperature reverses memory and anxiety-like behavior and reduced Aβ42 peptide levels in 3xTg-AD mice (Vandal et al., 2016). More recently, our group provided evidence that enhancement of thermogenesis through cold acclimation improves metabolic disorders and protects old 3xTg-AD mice from cold-induced tau phosphorylation (Tournissac et al., 2019). Supporting the link with age, cold-induced tau phosphorylation is potentiated in old mice compared to young mice (Tournissac et al., 2017). Altogether, these observations suggest that thermoregulatory mechanisms could be a potential therapeutic target in AD.

Beside thermoregulation, metabolic diseases share strong pathogenic links with AD. Indeed, induction of a diabetic phenotype such as glucose intolerance has been repeatedly shown to increase AD neuropathology in a mouse model of AD (Gratuze et al., 2017; Julien et al., 2010; Theriault et al., 2016; Vandal et al., 2014). Central insulin signaling defects and lower brain glucose metabolism are observed in AD (An et al., 2018; Arnold et al., 2018). It is estimated that one out of ten cases of AD is attributable to type 2 diabetes (T2D) (Biessels et al., 2006). These observations logically led to the idea of repurposing T2D drugs in AD (Yarchoan and Arnold, 2014). Insulin, thiazolidinediones and glucagon-like peptide-1 (GLP-1) analog are still the subject of clinical trials in dementia, albeit with mitigated results (Craft et al., 2012; Gejl et al., 2016; J. Liu et al., 2015). Thus, common metabolic targets between both diseases such as thermoregulatory defects are of interest to develop new therapeutic tools in AD.

Brown adipose tissue (BAT) is an essential thermogenic driver in mammals (Cannon and Nedergaard, 2004). The discovery of functional BAT in adults in 2009 has revived research on this tissue (Cypess et al., 2009; Virtanen et al., 2009). The ability of BAT thermogenesis to improve main metabolic disorders is now well-established in young (Hanssen et al., 2015; Ravussin et al., 2014; Schrauwen and van Marken Lichtenbelt, 2016) and old mice (Tournissac et al., 2019). Pharmacological tools have been developed in this direction. In particular, β3-adrenergic receptors (β3AR) agonists are being extensively used to stimulate β3AR located on brown adipocytes, thereby leading to lipolysis and UCP1 expression, the main marker of non-shivering thermogenesis (Arch, 2011; Nedergaard et al., 2001). CL-316,243 is a highly specific β3AR agonist frequently used in metabolic studies in rodents. It has been shown to improve blood glucose metabolism, insulin sensitivity, energy expenditure and to regulate lipids metabolism (Burkey et al., 2000; de Souza et al., 1997; Kim et al., 2006; Kumar et al., 2015; Labbé et al., 2016). Since β3AR agonists can correct metabolic disorders by enhancing BAT activity, they could tackle both T2D and AD at the same time. Importantly, β3AR agonists have been shown to stimulate BAT activity in humans (Baskin et al., 2018; Cypess et al., 2015) and one of these molecules, mirabegron (Myrbetriq^®^), is now approved for the treatment of overactive bladder (Chapple et al., 2014a). Therefore, β3AR agonists could rapidly be tested in humans for dementia.

We hypothesized that pharmacological stimulation of BAT thermogenesis through β3AR agonist treatment could curtail AD neuropathology and improve memory as well as correcting thermoregulatory and metabolic deficits. To verify this hypothesis, 15-month-old non-transgenic (NonTg) and 3xTg-AD mice received daily CL-316,243 (1 mg/kg) or saline injections for a month.

## MATERIAL AND METHODS

### Animals

The triple-transgenic mouse model of AD (homozygous 3xTg-AD; APP_swe_, PS1_M146V_, tau_P301L_) developing both amyloid and tau pathologies in the brain with age was used here (Oddo et al., 2003). We selected 15-month-old 3xTg-AD and NonTg controls, at an age when 3xTg-AD mice have extended plaques and tangles in the brain, as well as cognitive deficits (Belfiore et al., 2018; Bories et al., 2012; St-Amour et al., 2014; Vandal et al., 2015). Animals were produced at our animal facility and all maintained in the same genetic background (C57BL6/129SvJ) by backcrossing every 8-10 generations. Forty-two (42) mice were used for all experiments (n= 9-12 mice per group) and 9 mice were added for behavioral and glucose tolerance tests for a total of 51 mice (n= 9-16 mice per group).

Mice were housed one to five per cage at a housing temperature of 23 °C, with a 12:12 hours light-dark cycle (light phase from 7 a.m. to 7 p.m). Animals had ad libitum access to water and chow (Teklad 2018, Harlan Laboratories, Canada). Only males were used here to avoid temperature variation induced by the estrous cycle of female mice (Weinert et al., 2004). Food consumption was evaluated by weighing the diet of each cage and averaged for each mouse per day per cage every four days during the one-month treatment, and three weeks before the beginning of the experiment. At the end of the experiment, all mice were put under deep anesthesia with ketamine/xylazine intraperitoneal (i.p.) injection (100 mg/kg ketamine, 10 mg/kg xylazine) and immediately placed under a heating pad to maintain body temperature until complete loss of posterior paw reflex. Then, mice were rapidly sacrificed by intracardiac perfusion with 0.1 M phosphate buffer saline (PBS) solution containing phosphatases (sodium pyrophosphate, 1 mM and sodium fluoride, 50 mM) and proteases (Sigmafast protease inhibitor tablets, Sigma-Aldrich, St-Louis, USA) inhibitors. All experiments were performed in accordance with the Canadian Council on Animal Care and were approved by the Institutional Committee of the Centre Hospitalier de l’Université Laval (CHUL).

### CL-316,243 treatment

CL-316,243 was selected to stimulate BAT thermogenesis because it is one of the most selective β3AR agonists in rodents (β1: β2: β3 = 0:1:100,000) and its safety and efficacy has been confirmed in multiple studies (Bloom et al., 1992; Caron et al., 2017; Danysz et al., 2018; Ghorbani et al., 2012; Yoshida et al., 1994). Two to three weeks of daily injection at a dose of 1 mg/kg per day are necessary to improve metabolic disorders (Burkey et al., 2000; de Souza et al., 1997; Kim et al., 2006; Kumar et al., 2015; Labbé et al., 2016). Thus, mice were injected i.p. every day for a month with a weight-adjusted dose of CL-316,243 (1 mg/kg) or an equivalent volume of saline (the vehicle) at the same hour of the day (4 p.m.) from 15 to 16 months of age (Fig. 1A). Mice were weighed every day before each i.p. injection. Mice were sacrificed the morning after the last injection.

**Figure 1.**
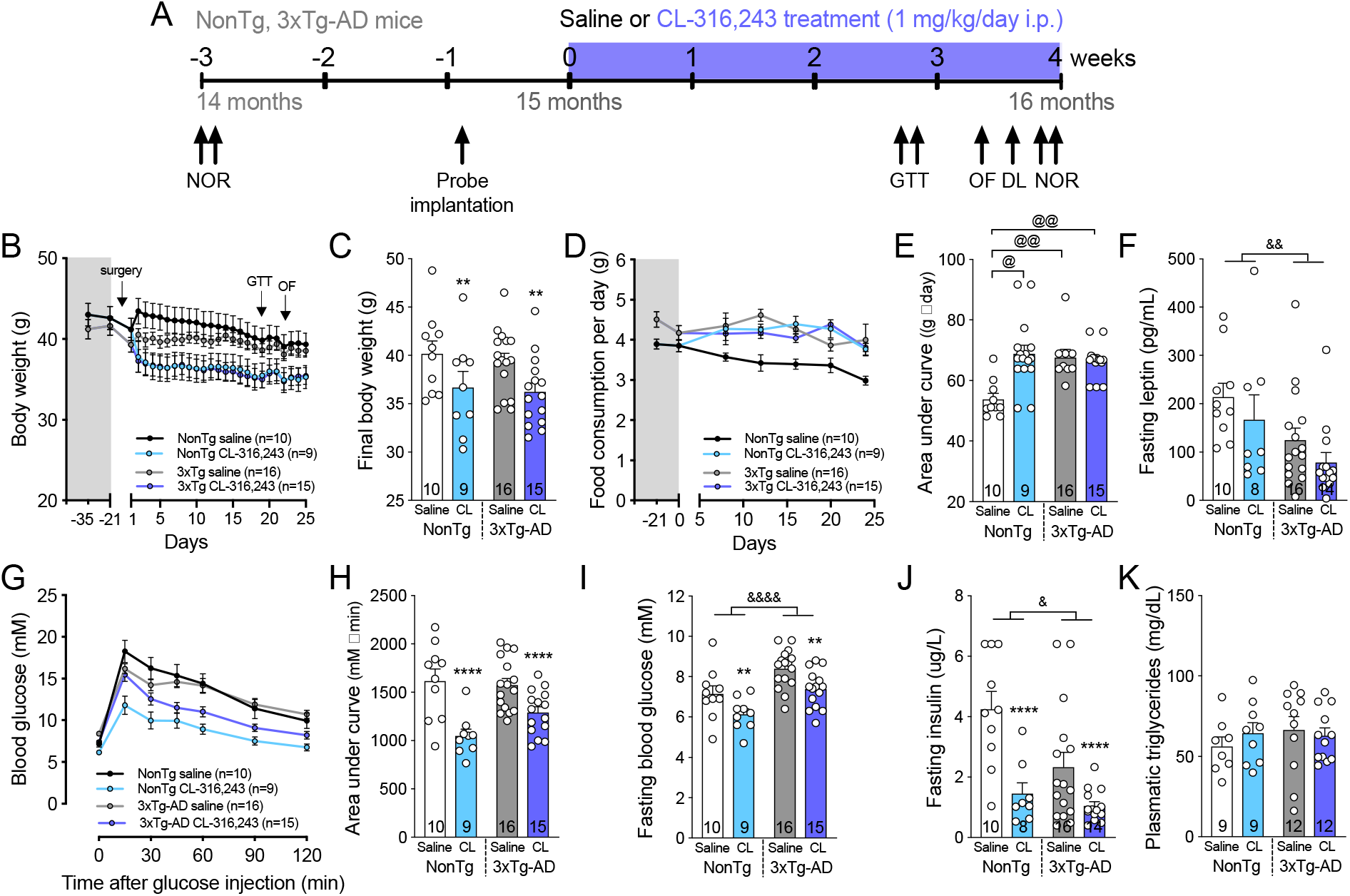
β3AR stimulation improves peripheral glucose metabolism in NonTg and 3xTg-AD mice. A: Schematic description of the experimentation. B: Body weight before and during the one-month experiment and C: at the end of the experiment. D: Food consumed per day per mice three weeks before and over the one-month treatment and E: area under curve of food consumption over the one-month treatment. F: Leptin measured in plasma of fasted mice. G: GTT performed after 3 weeks of experiment and H: area under curve of the GTT. I: Fasting blood glucose and J: plasmatic insulin measured during the GTT. I: Triglycerides measured in the plasma sampled at the end of the experiment. Data are represented as mean ± SEM (n/group indicated in graphics). Statistics: Two-way ANOVA, effect of treatment: **p<0.01; ****p<0.0001; effect of genotype: ^&^p<0.05; ^&&^p<0.01; ^&&&&^p<0.0001 (C,F,H,I,J). Kruskal-Wallis, Dunn’s post-hoc test: ^@^p<0.05; ^@@^p<0.01 (E). Abbreviations: 3xTg-AD: triple transgenic mice; DL: dark-light box test; GTT: glucose tolerance test; NonTg: non-transgenic mice; NOR: novel object recognition test; OF: open field.

### Body temperature measurement and analysis

Telemetric probes (Anipill, Caen, France) were used to record body temperature of the animals every hour during the one-month experiment without manipulation. Probes were implanted in the intraperitoneal cavity under isoflurane anesthesia a week before the beginning of the treatment to allow recovery from the surgery (Fig. 1A). Heat pads were used throughout the procedure to avoid hypothermia. The time under anesthesia was similar between mice and lasted approximately 10 minutes. Then, the animals were kept under heat pads during the waking period. Body temperature was analyzed during the two first weeks of treatment, before animals underwent glucose tolerance and behavioral tests, to avoid resulting interference in circadian rhythms.

In order to assess potential phase advances or delays and to visualize endogenous rhythmicity and regularity of the mice circadian clock, Clocklab software (Actimetrics Inc., Evanston, Illinois, USA) provided the further information: individual mean daily offsets in hours (determined as the time of the first six successive bins when temperature was lower than the mean diurnal temperature), offset standard deviation in hours and mean duration of a total temperature cycle in hours. We performed our analyses based on daily offsets (the end of the active period of the mice) because daily injections of the drug were performed only three hours before the lights turned off, thus influencing the onsets (the beginning of the active period). Other following parameters were calculated for each individual: mean body temperature in °C during the dark (from 7 p.m. to 7 a.m.) and light phase (from 7 a.m. to 7 p.m.) and mean amplitude of temperature (T°max-T°min) in °C during the dark and light phase.

### Glucose tolerance test, fasting blood glucose, leptin and insulin

Glucose tolerance test (GTT) was performed at the end of the third week of treatment (Fig. 1A) (Vandal et al., 2015). Mice were fasted for 6 hours (from 8 a.m. to 2 p.m.). Then, glucose was injected i.p. at 1 g/kg and blood glucose was measured regularly during 2 hours with a glucometer (OneTouch UltraMini; LifeScan, Milpitas, CA) in a blood drop sampled from the saphenous vein. Leptin and insulin were assessed by ELISAs (Leptin Mouse ELISA kit, ab100718, Abcam; Mouse insulin ELISA, #10-1247-01, Mercodia, Sweden) following manufacturer’s instructions in plasma sampled in the saphenous vein after the 6-hour fasting during the GTT test.

### Behavioral tests

Behavioral tests were performed during the fourth week of treatment with a recovery time of at least 24 hours between tests (Fig. 1A). The novel object recognition (NOR) test was also performed three weeks before the beginning of the treatment to obtain a baseline index ratio for each animal (see below). Mice were acclimated overnight to the testing room located next to the housing room.

Locomotor activity was assessed with the open field test (Dal-Pan et al., 2016). Mice were placed in a 40 cm × 40 cm × 40 cm translucent Plexiglas box for an hour. Movements were tracked with photobeam breaks (San Diego Instruments). The total distance traveled (voluntary horizontal movement) and the average speed were compared between groups.

Anxiety behavior was evaluated with the dark-light box test (Vandal et al., 2016). Mice were put in the center of the dark compartment with an opening to the light compartment. The time spent in the light compartment and the latency to do the first exploration (nose latency) of the light compartment were measured during a 5-minute trial.

Memory deficits were evaluated with the NOR test. That test detects behavioral deficits from 12 months in 3xTg-AD mice and is one of the less stressful behavioral test (Arsenault et al., 2011; Clinton et al., 2007; St-Amour et al., 2014; Vandal et al., 2016). It evaluates recognition memory and corresponds to episodic memory that is early affected in AD (Antunes and Biala, 2012; Leger et al., 2013; Wolf et al., 2016). Mice were first placed in a 29.2 cm × 19 cm × 12.7 cm cage with two identical objects for 5 minutes during the acquisition phase. After an hour in their housing cage, mice returned in the testing cage containing a familiar and a novel object for the test phase. Recognition index (RI) corresponds to the time spent exploring the novel object divided by the total time of exploration during the test phase multiplied by 100. A 50% RI corresponds to an equal exploration between the novel and the familiar object. Mice exploring less than 6 seconds each object during the acquisition phase or less than 4 seconds during the test phase were excluded from the RI analysis. Mice were assigned to the treated or the control group at 15 months of age with caution to homogenize memory performance (baseline RI) between groups at the beginning of the experiment.

### Tissue preparation for postmortem analysis

Intracardiac blood sampled just before intracardiac perfusion in a heparinized tub was centrifuged at 3000 rpm for 5 minutes, and resulting plasma kept frozen at −80 °C until analysis. The first hemisphere and interscapular BAT were rapidly dissected and frozen at −80 °C until processing. The second hemisphere was either fixed in 4% paraformaldehyde for 48 hours and transferred in a 20% sucrose solution until sectioning (3-4 mice per group) or frozen and kept at −80 °C.

### Protein extractions

For the hippocampus, frozen samples were homogenized in 8 volumes of a lysis buffer (150 mM NaCl, 10 mM NaH2PO4, 0.5% sodium deoxycholate, 0.5% sodium dodecyl sulfate, 1% Triton X-100) containing a cocktail of protease and phosphatase inhibitors (Bimake, Houston, TX), sonicated (3 × 45 s in a Sonic Dismembrator apparatus, Thermo Fisher Scientific, Waltham, MA) and centrifuged (100,000g, 20 min, 4°C), resulting in a detergent-soluble fraction (cytosolic, extracellular and membrane-bound proteins). The remaining pellets from ultracentrifugation were resuspended in formic acid, resulting in a detergent-insoluble fraction (insoluble proteins fraction). The resultant suspension was sonicated and centrifuged (13,000g, 20 min, 4 °C), acid formic was evaporated and proteins were either solubilized in Laemmli’s buffer for Western blot or in a 5 M guanidium solution in Tris-HCl 50 mM for Aβ peptides ELISAs as previously described (Tremblay et al., 2017). Proteins from the BAT were extracted in the lysis buffer only. Protein concentrations were evaluated with a bicinchoninic acid assay (BCA, Pierce, Rockford, IL, USA).

### Western immunoblotting

15 μg and 10 μg of proteins of hippocampus and BAT homogenates, respectively, were loaded and separated by SDS-PAGE, as previously described (Vandal et al., 2014). The list of antibodies used in this study is available in Table S2. Homogenates were all run on the same gel for each experiment.

### Aβ40 and Aβ42 peptides quantification

Aβ peptides were quantified in protein extracts from the hippocampus. Aβ40 and Aβ42 were measured in detergent-soluble and detergent-insoluble fractions using a human β-amyloid ELISA (Wako, Osaka, Japan) according to the manufacturer’s instructions. Plates were read at 450 nm using a Synergy™ HT multi-detection microplate reader (Biotek, Winooski, VT).

### High-performance liquid chromatography (HPLC)

HPLC was used to measure the level of norepinephrine in BAT. An average of 10 mg of BAT was homogenized in perchloric acid (0.1 N) and centrifuged 10 minutes at 4 °C at 13 000 rpm. 5 μL of the supernatant was injected in the HPLC with electrochemical detection (Water 717 plus Autosampler automatic injector, Waters 1525 Binary Pump) as previously described (Bousquet et al., 2011).

### Statistical analysis

Data are represented as means ± standard error of the mean (SEM). Statistical analysis and the number of samples per group are specified in each figure and legend. Bartlett’s tests were used to rule out the inequality of variances between the groups. Two-way ANOVA (two independent variables: genotype and treatment) was used in case of equal variances. Repeated measures two-way ANOVA was executed to compare recurrent measurements in same animals. Tukey’s test was used for post-hoc analysis. In case of unequal variances, a Kruskal-Wallis followed by a Dunn’s post-hoc test was performed. An unpaired Student’s t-test was performed when only two groups were compared, with a Welch correction in case of unequal variances. Paired t-test was executed for the before-and-after on comparison on the same animals. One sample t-test was used to compare means to a theoretical value (for the NOR test). Correlations between variables were investigated using linear regression analyses. All statistical analyses were performed with Prism 7 (GraphPad Software Inc., San Diego, CA, USA) or JMP (version 13.2.0; SAS Institute Inc., Cary, IL, USA) software and statistical significance was set at p<0.05.

## RESULTS

### 1. β3AR stimulation improves peripheral glucose metabolism in old mice

NonTg and 3xTg-AD mice received CL-316,243 or saline i.p. at a dose of 1 mg/kg every day for a month from 15 to 16 months of age (Fig. 1A). To verify whether CL-316,243 affects energy balance, mice were weighed every day before each i.p. injection and food consumption was evaluated three weeks before the beginning of the experiment and then, every four days during the one-month treatment. First, we found that CL-316,243 injections induced persisting weight loss in both NonTg and 3xTg-AD mice (Fig. 1B,C). At the beginning of the experiment, food consumption recorded in the previous 21 days was higher in 3xTg-AD compared to NonTg mice (average of 4.7 g/day/mice for 3xTg-AD versus 3.9 g/day/mice for NonTg mice; unpaired t-test with Welch’s correction: p=0.0105) (Fig. 1D), consistent with lower plasmatic leptin in fasted 3xTg-AD mice compared to NonTg (Fig. 1F). Over the one-month period, β3AR stimulation increased food consumption in NonTg mice up to levels of transgenic mice (Fig. 1D,E). The GTT revealed that CL-316,243-treated mice displayed a stronger control over glucose levels compared to control, independently of the genotype (Fig. 1G,H). Fasting blood glucose and insulin were also lower following three weeks of CL-316,243 administration (Fig. 1I,J). However, levels of plasmatic triglycerides were unchanged (Fig. 1K). Overall, our data indicate that β3AR stimulation led to an improved pattern of metabolic determinants in the periphery in both NonTg and 3xTg-AD mice at 15-16 months of age.

### 2. β3AR stimulation increases brown adipose tissue thermogenesis

A telemetric probe implanted a week before the beginning of the treatment revealed daily variation in body temperature corresponding to the sleep-wake cycles of mice (Fig. 2A, Fig. S1). The mean amplitude of body temperature was larger by 0.4 °C in 3xTg-AD than in NonTg mice during both 12-h light and 12-h dark phases (Fig. 2A-C, Fig. S1D), in agreement with thermoregulation defects previously reported in the same model (Vandal et al., 2016). CL-316,243 treatment further increased the amplitude of body temperature during the light phase (from 7 a.m. to 7 p.m.) compared to saline injections (Fig. 2B), corresponding to the injection time, but not during the dark phase (Fig. 2C). This is consistent with higher area under curve of body temperature measured after the first injection of CL-316,243 (between 4 p.m. and 12 p.m.) (Fig. 2D,E).

**Figure 2.**
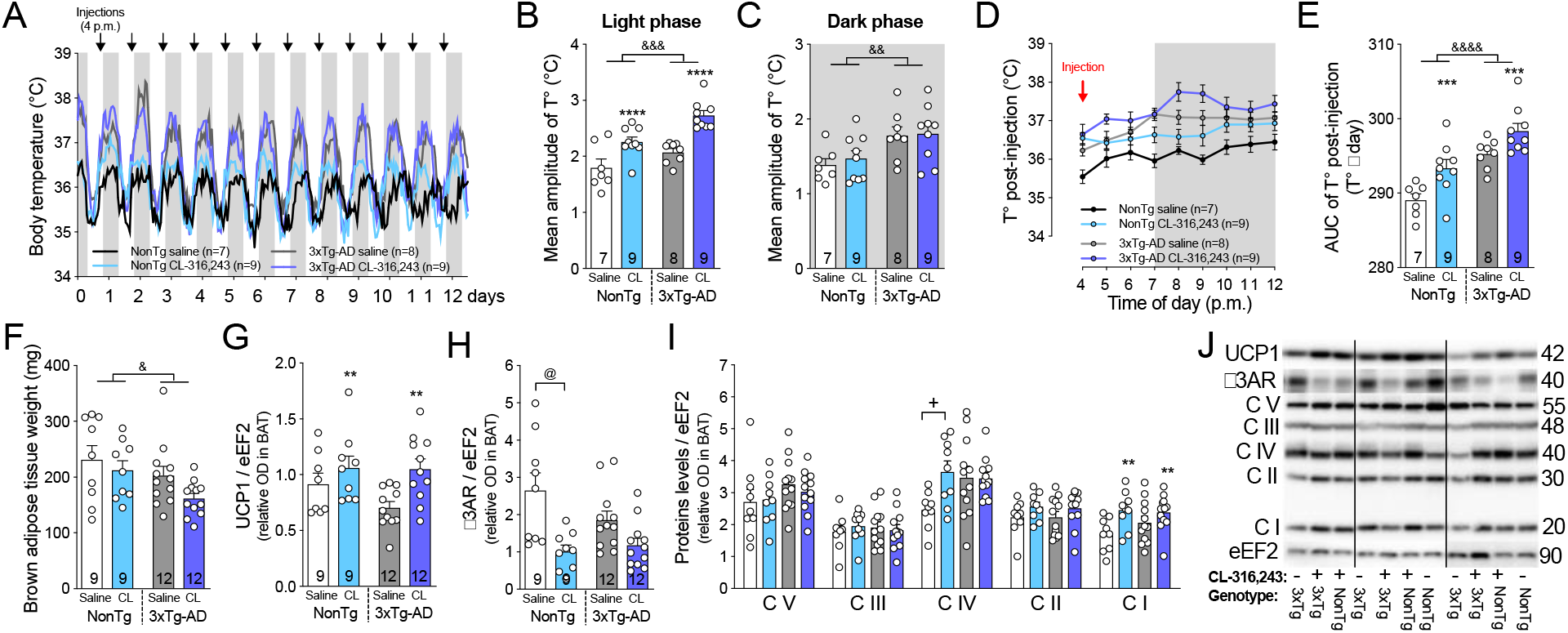
CL-316,243 treatment increases brown adipose tissue thermogenesis. A: Graphical representation of body temperature recorded hourly by telemetric probe in the first two weeks of experiment (before glucose tolerance and behavioral tests). B: Mean amplitude of body temperature during the light (12-h, from 7 a.m.) and C: the dark phase (12-h, from 7 p.m.). D: Body temperature (T°) and E: area under curve of the T° after the first CL-316,243 or saline i.p. injection. F: Interscapular BAT weights. Levels of G: UCP1, H: β3AR and I: mitochondrial oxidative phosphorylation system on eEF2 proteins measured in BAT by Western Blot. J: Examples of Western Blots. Homogenates were all run on the same gel, but consecutive bands were not taken for all representative photo examples. Data are represented as mean ± SEM (n/group indicated in each column). Statistics: Two-way ANOVA, effect of treatment: **p<0.01; ***p<0.001; ****p<0.0001; effect of genotype: ^&^p<0.05; ^&&^p<0.01; ^&&&^p<0.001; ^&&&&^p<0.0001 (B,C,E-G,I); Tukey’s post-hoc test: ^+^p<0.05. Kruskal-Wallis (I), Dunn’s post-hoc test: ^@^ p<0.05 (H). Abbreviations: AUC: area under curve; 3xTg-AD: triple transgenic mice; β3AR: β3 adrenergic receptor; BAT: brown adipose tissue; CV, CIII, CIV, CII, CI: mitochondrial oxidative phosphorylation system complex V, III, IV, II, I; eEF2: eukaryotic elongation factor 2; NonTg: non-transgenic mice; OD: optical density; T°: temperature; UCP1: uncoupling protein 1.

Since CL-316,243 is well known to improve thermogenesis capacity in mice (Labbé et al., 2016; Poher et al., 2015; Xiao et al., 2015), we then verified whether it was also effective in old NonTg and 3xTg-AD mice. First, interscapular BAT weight was slightly lower in 3xTg-AD compared to NonTg mice, but was not affected by the treatment (Fig. 2F). However, CL-316,243 administration increased the level of UCP1 protein in the BAT of both genotypes (Fig. 2G) but did not affect the norepinephrine content (Fig. S1E). Further confirming that CL-316,243 interacted with β3AR, levels of β3AR in BAT were significantly decreased only in NonTg treated mice, despite a tendency also in 3xTg-AD mice (Fig. 2H). We then measured complexes I to V of the mitochondrial oxidative phosphorylation complex that are involved in heat production during thermogenesis in BAT (Nam and Cooper, 2015). Complex I was increased in NonTg and 3xTg-AD mice following CL-316,243 administration (Fig. 2I,J). CL-316,243 increased complex IV in NonTg, but not in 3xTg-AD mice, whereas complexes II, III and V remained unchanged in both models. Altogether, our data show that the β3AR agonist administration improves BAT thermogenesis and heat production in 16-month-old mice.

### 3. β3AR stimulation reverses memory deficits in 16-month-old 3xTg-AD mice

To determine whether CL-316,243 treatment exerted cognitive benefits in the 3xTg-AD mouse, recognition memory was evaluated with the NOR test 3 weeks before the beginning of the treatment (baseline, 14-month-old) and after the one-month treatment (final, 16-month-old) (Fig. 1A). The NOR test was selected because of its sensitivity and reliability to detect memory deficits in the 3xTg-AD mice at 12 months and older (Fig. 3A) (Arsenault et al., 2011; Clinton et al., 2007; Vandal et al., 2016). Comparing RI before (14 months) and after (16 months) the treatment revealed that one-month treatment with CL-316,243 increased by 19% the ability to recognize the new object in 3xTg-AD mice (paired t-test: p=0.0041), while the change in RI was not significantly different in NonTg or saline-injected 3xTg-AD mice (Fig. 3B,C). CL-316,243 from 15 to 16 months improved memory recognition in 3xTg-AD mice (RI=65% in CL-316,243-injected mice, one sample t-test versus 50%: p=0.0013), but not in NonTg mice (Fig. 3D). The improved RI in 3xTg-AD treated mice was confirmed by the higher time spent on the novel (N) versus the familial (O) object (Fig. 3E). These differences were not explained by changes in exploratory behavior, as the mean duration of exploration was similar between groups (Fig. 3F). Finally, the percent change in RI before and after the treatment was positively correlated with UCP1 levels in BAT in 3xTg-AD (r^2^=0.37) but not in NonTg mice, suggesting a link between improved thermogenesis and memory (Fig. 3G).

**Figure 3.**
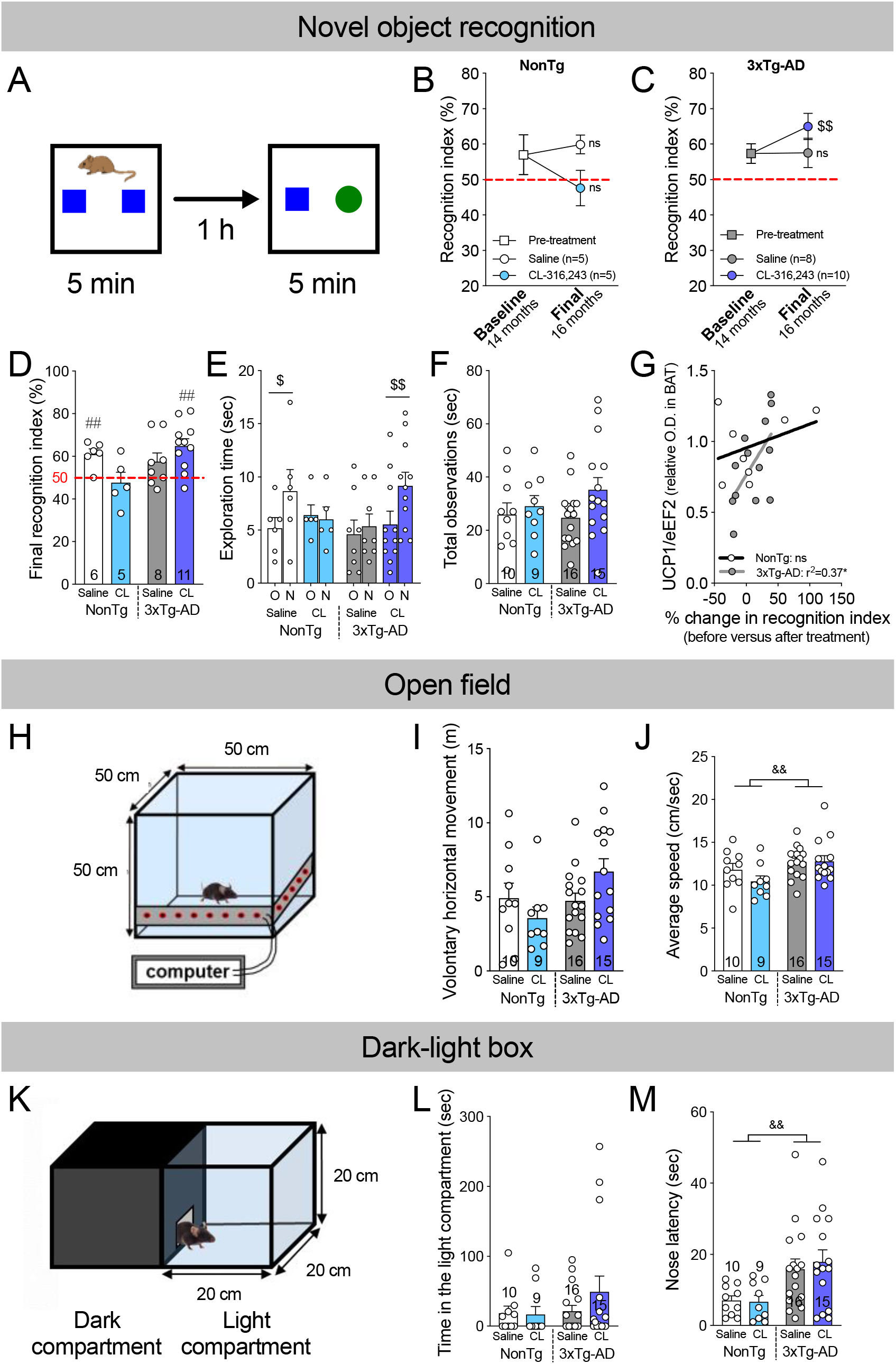
β3AR stimulation reverses memory deficits in 16-month-old 3xTg-AD. A: Description of the novel object recognition test. Recognition indexes (RI) assessed before and after the treatment with CL-316,243 or saline in B: NonTg and C: 3xTg-AD mice. D: Final recognition index, E: time spent exploring the old (O) and the novel (N) object during the 5-minutes acquisition phase and F: total observations measured at the end of the experiment (final, 16-month-old mice). G: Correlation between % change in recognition index before and after the treatment and UCP1 levels measured in BAT. H: Representation of the open field apparatus. I: Total distance traveled and J: average speed during the one-hour test. K: Representation of the dark-light box. L: Time spent in the light compartment and M: latency to do the first exploration in the light compartment. Recognition index = (time exploring the novel object / total exploration time) × 100. Data are represented as mean ± SEM (n/group indicated in graphics). Statistics: Paired t-test (Baseline versus final recognition index (B,C); Old (O) versus novel (N) object (E)): ^$^p<0.05; ^$$^p<0.01 (B,C,E). One sample t-test versus 50% (random chance): ^##^p<0.01 (D). Pearson r correlation: *p<0.05 (G). Two-way ANOVA, effect of genotype: ^&&^p<0.01 (J,M). Abbreviations: 3xTg-AD: triple transgenic mice; eEF2: eukaryotic elongation factor 2; O.D., optical density; NonTg: non-transgenic mice; UCP1: uncoupling protein 1.

We then verified that locomotor activity was not affected by CL-316,243 injections, as showed by comparable distance traveled and average speed of the mice during the open field test (Fig. 3H-J). Nonetheless, 2-way ANOVA revealed that 3xTg-AD mice displayed a higher average speed during the one-hour session compared to NonTg mice (Fig. 3J).

Anxiety is frequently observed in AD patients and is replicated in 3xTg-AD mice (Hebda-Bauer et al., 2013; St-Amour et al., 2014; Vandal et al., 2016). The time spent in the illuminated compartment was not significantly different between groups in the present cohorts of animals (Fig. 3L). However, 3xTg-AD mice delayed their first exploration in the light chamber, as measured by the latency of the first nose entry in the light compartment (nose poke latency) (Fig. 3M), corroborating an anxiety-like behavior in this model.

Overall, one-month administration of CL-316,243 improved recognition memory assessed at 16 months in 3xTg-AD mice, without affecting locomotion nor anxiety-like behavior.

### 4. β3AR stimulation reduces insoluble Aβ42/Aβ40 ratio in the hippocampus of 3xTg-AD mice

The main neuropathological markers of AD, amyloid plaques and tau pathology (Iqbal et al., 2016; Selkoe and Hardy, 2016), progressively develop in the brain of 3xTg-AD mice (Belfiore et al., 2018; Oddo et al., 2003; Vandal et al., 2015). Although total Aβ42 and Aβ40 peptides in either soluble or insoluble fractions remained unchanged by the treatment in the hippocampus (Fig. 4A-D), we observed a 27% decrease in insoluble Aβ42/Aβ40 ratio in CL-316,243-treated mice compared to saline-injected 3xTg-AD mice (Fig. 4E). We subsequently assessed the effect of the β3AR agonist on proteins implicated in the production (beta-secretase 1 (BACE-1), amyloid precursor protein (APP), APP C-terminal, sAPPα) and clearance or degradation (low density lipoprotein receptor related protein 1 (LRP1), receptor of advanced glycation end products (RAGE), insulin degrading enzyme (IDE), X11α) of Aβ peptides (Table S1) (Donahue et al., 2006; Farris et al., 2003; Selkoe and Hardy, 2016). Levels of BACE-1, IDE, LRP1 and RAGE in the hippocampus did not differ between groups, whereas those of X11α were lower in 3xTg-AD mice compared to NonTg mice (Table S1) as observed in the brain of AD individuals (Tremblay et al., 2017).

**Figure 4.**
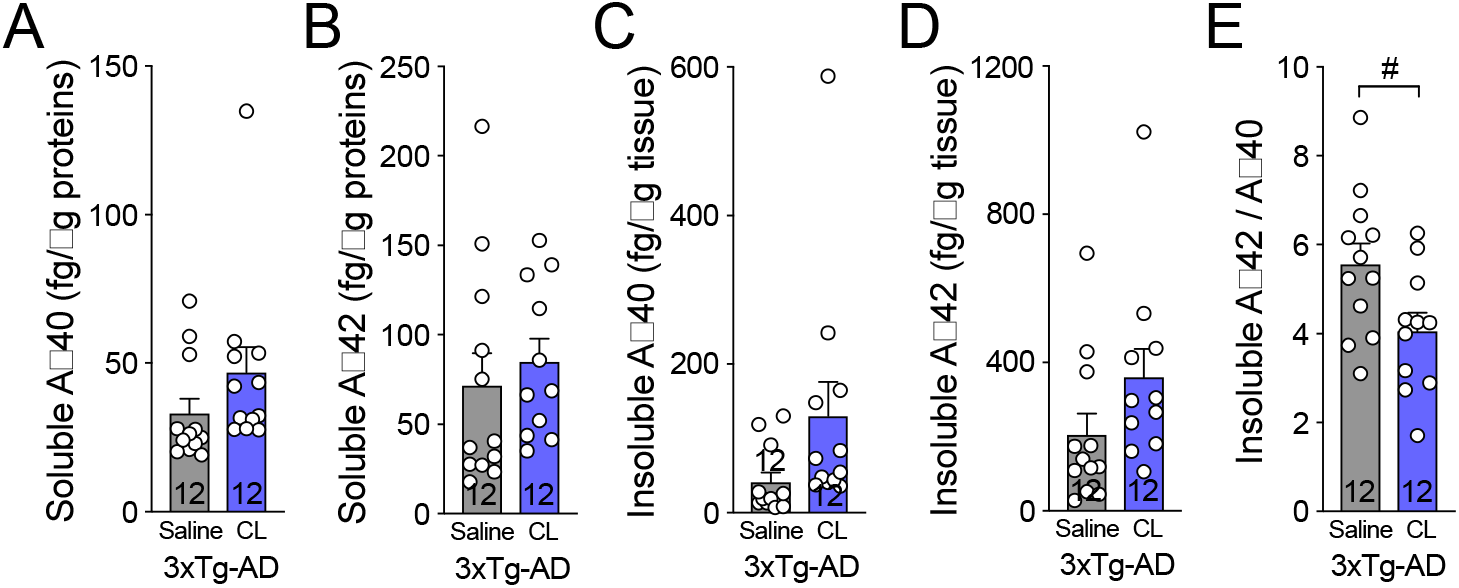
β3AR stimulation reduces insoluble Aβ42/Aβ40 ratio in the hippocampus of 3xTg-AD mice. Human Aβ40 and Aβ42 peptides measured by ELISA in the A,B: detergent-soluble and in the C,D: detergent-insoluble fractions of hippocampus homogenates of 3xTg-AD mice, respectively. E: Ratio of insoluble Aβ42 on Aβ40 peptides in the hippocampus. Data are represented as mean ± SEM (n/group indicated in bars). Statistics: Unpaired Student t-test:^#^p<0.05 (A-E). Abbreviations: 3xTg-AD: triple transgenic mice; Aβ: amyloid-beta; NonTg: non-transgenic mice.

We then assessed the level of phosphorylated and total tau protein in the detergent-soluble (cytosolic and membrane proteins) and detergent-insoluble (aggregated proteins) fractions of hippocampus homogenates by Western Blot. We did not find any significant effect of CL-316,243 (Fig. 5) but confirmed that the 3xTg-AD mice display higher total and hyperphosphorylated tau proteins compared to NonTg mice. Main kinases involved in tau phosphorylation (glycogen synthase kinase 3 β (GSK3β) and protein kinase B known as AKT) were also unchanged (Table S1).

**Figure 5.**
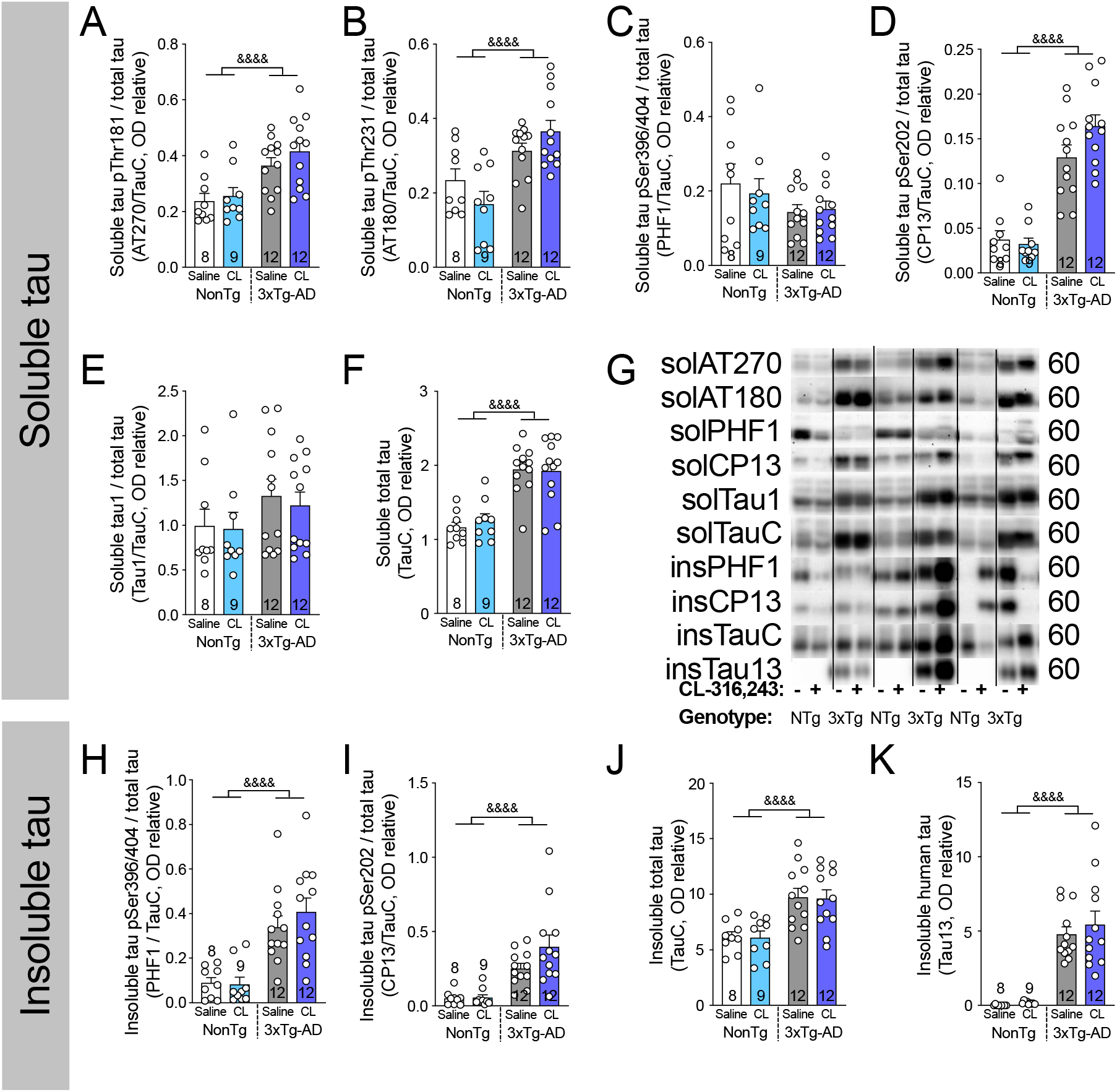
β3AR stimulation has no effect on tau phosphorylation in the hippocampus of 3xTg-AD mice. Phosphorylated and total tau protein measured in A-F: detergent-soluble and H-K: detergent-insoluble fractions of hippocampus homogenates of non-transgenic and 3xTg-AD mice. G: Examples of Western Blots. Homogenates were all run on the same gel, but consecutive bands were not taken for all representative photo examples and were cut-pasted to match the histogram order. Data are represented as mean ± SEM (n/group indicated in bars). Statistics: Two-way ANOVA, effect of genotype: ^&&&&^p<0.0001 (A-K). Abbreviations: 3xTg-AD: triple transgenic mice; ins: insoluble; NonTg: non-transgenic mice; OD: optical density; sol: soluble.

Synaptic deficits are one of the earliest markers of AD, correlating with symptoms (Arendt, 2009; Tremblay et al., 2017). The levels of synaptic proteins were not modified by the treatment (Table S1). However, drebrin protein was decreased specifically in 3xTg-AD mice (Two-way ANOVA, effect of genotype: p=0.0045), as previously showed in the brain of AD subjects (Calon et al., 2004; Julien et al., 2008).

Since glucose transporters and uptake are decreased in AD (An et al., 2018; Kalaria and Harik, 1989; Y. Liu et al., 2008; Simpson et al., 1994), we assessed glucose transporter 1 (GLUT1) levels in the hippocampus of the mice. While we did not detect any effect of the CL-316,243 treatment, GLUT1 levels were decreased in the hippocampus of 3xTg-AD mice compared to NonTg mice, at both the endothelial (50 kDa) and astrocytic (45 kDa) isoforms (Table S1), corroborating defects in glucose uptake, changes in blood-brain barrier transporters and decreased cerebral vascular volume observed in this mouse model of AD (Bourasset et al., 2009; T. M. Do et al., 2014; 2016; Nicholson et al., 2010).

## DISCUSSION

The present study aimed at investigating whether β3AR agonist administration induces BAT thermogenesis and exerts an effect on cognitive behavior and AD neuropathology in a mouse model of the disease. The 3xTg-AD mice was selected to test the effect of β3AR stimulation in AD because this model displays age-dependent metabolic and thermoregulatory deficits and was shown to respond to thermoneutrality and BAT stimulation induced by repeated cold exposure (Knight et al., 2013; Tournissac et al., 2019; Vandal et al., 2016). This led to the hypothesis that pharmacological BAT stimulation could exert benefit on AD-like behavior and neuropathology. We thus treated 15-month-old NonTg and 3xTg-AD mice with the selective β3AR agonist CL-316,243 or saline for a month. We found that chronic administration of the agonist stimulated BAT thermogenesis and improved glucose homeostasis. Enhanced thermogenesis was associated with improved recognition memory in 16 months 3xTg-AD mice and reduced insoluble Aβ42/Aβ40 ratio in the hippocampus, while tau pathology remained unaffected.

### 1. β3AR agonist: a two in one strategy to target both metabolic and thermoregulatory defects

The most well-known characteristic of CL-316,243 is to improve metabolic disorders through enhanced BAT activity (Burkey et al., 2000; de Souza et al., 1997; Kim et al., 2006; Kumar et al., 2015; Labbé et al., 2016) (Cypess et al., 2015; Loh et al., 2018). Whether this effect is maintained with age remains unknown. UCP1 and NADH dehydrogenase 1 beta subcomplex subunit 8 (NDUFB8 or Complex I) are both recognized markers of BAT activation (Nam and Cooper, 2015). Higher levels of UCP1 and Complex I in the BAT detected here indicate a sustained effect of CL-316,243 on BAT thermogenesis after a one-month chronic treatment in old mice. This is consistent with strong previous evidence that β3AR stimulation leads to increased thermogenic activity of BAT in young mice (Labbé et al., 2016; Poher et al., 2015; Xiao et al., 2015) and humans as well (Cypess et al., 2015; Loh et al., 2018). It also shows that chronic β3AR administration does not induce significant receptor desensitization, despite decreased levels of BAT β3AR following treatment, in agreement with previous works (Nedergaard and Cannon, 2013). Overall, our data now confirms that β3AR-induced thermogenesis is still effective in a 15-month-old mouse, following a one-month treatment with CL-316,243, regardless of the presence of AD neuropathology.

In the present study, β3AR agonist induced weight loss in both NonTg and 3xTg-AD mice. Food consumption was higher in 3xTg-AD compared to NonTg mice, an observation previously reported in this model (Adebakin et al., 2012; K. Do et al., 2018), and CL-316,243 increased food intake in NonTg mice. These data are consistent with increased energetic expenditure compensated with higher calorie intake following β3AR stimulation (Gavrilova et al., 2000; Xiao et al., 2015). We previously showed that female 3xTg-AD mice display age-dependent glucose intolerance starting from 12 months (Vandal et al., 2015). While the difference in glucose tolerance between NonTg and 3xTg-AD mice was not frank in the present work considering that only males were used, fasting blood glucose in 3xTg-AD was higher than in NonTg mice. It is noteworthy that the CL-316,243 improved peripheral glucose metabolism and fasted insulin in 16-month-old mice of both genotypes, suggesting an effect independent of AD neuropathology. Of note, the improvement was observed in mice fed with a diet not expected to induce metabolic defects, suggesting benefits even in non-diabetic animals. This is in line with a previous study showing improved glucose metabolism with CL-316,243 administration even in mice displaying a normal response to glucose (Xiao et al., 2015).

Using hourly telemetric recordings of body temperature, we noted that 3xTg-AD mice displayed wider amplitudes of body temperature throughout the day compared to NonTg mice, corroborating previous findings showing that AD neuropathology impacts thermoregulation (Vandal et al., 2016). On the other hand, β3AR agonist increased amplitude of body temperature during the light phase, corresponding to the period of drug administration, but not during the dark phase. The wider amplitude in body temperature is consistent with higher body temperature after CL-316,243 injection, as shown by the temperature recorded few hours after the first i.p. injection and a previous work (Szentirmai and Kapás, 2017). However, we did not observed chronic hyperthermia, which would have been a major side effect of β3AR agonists, perhaps compromising potential translation to clinical use.

### 2. β3AR stimulation reverses memory deficits in old 3xTg-AD mice

An important result of our study is the reversal of recognition memory deficit induced by β3AR stimulation in 16-month-old 3xTg-AD mice. Indeed, we observed a 19% increase of RI between baseline (14 months) and post-treatment evaluation (16 months). These results were not explained by changes in exploratory behavior or locomotor activity. The RI of NonTg mice following treatment did not reach statistical significance perhaps due to lower statistical power (n=5), but was significantly different from 50% in saline-injected mice. Nevertheless, the data also suggest that β3AR-induced improvement in memory was specific to 3xTg-AD mice, possibly through an effect related to AD neuropathology. The randomized-start design ensured that all animals were similar before undergoing saline or CL-316,243 treatment. One study also found memory improvement following CL-316,243 administration in Aβ-injected chicks (Gibbs et al., 2010), but no previous report in mice was found in the literature. Thus, our results are consistent with a disease-modifying effect of β3AR stimulation in the 3xTg-AD mice.

Since metabolic disorders also alter cognitive function and lead to memory defects (Abbondante et al., 2014; Gunstad et al., 2010; Rajasekar et al., 2017; Takeda et al., 2010; Tong et al., 2019), improved peripheral metabolism could be involved in better recognition memory in 3xTg-AD mice. However, it is not excluded that CL-316,243 has a direct effect in the CNS. Indeed, β3AR are present in various regions of the brain, although to a much lower extent compared to the BAT (Evans et al., 1996; Summers et al., 1995), but their physiological roles in the central nervous system (CNS) are not known. While it has been shown that CL-316,243 increases sleep duration in mice (Szentirmai and Kapás, 2017) and reduces Aβ-induced long-term memory deficits in chicks (Gibbs et al., 2010), the behavioral effects of β3AR agonists has not been the subject of intense investigation. Nonetheless, another β3AR agonist, SR856611A (Amibegron^®^), has been shown to improve anxiety and depressive-like symptoms in rodents, as evaluated by the forced swim test and the elevated plus maze (Consoli et al., 2007; Stemmelin et al., 2008; Tamburella et al., 2010), and has been the subject of phase III clinical trial for depression (NCT00252330). In the present work, we did not detect changes in anxiety-like behavior with the dark-light box emergence test following CL-316,243 administration. What stand up from our data is that enhanced UCP1 levels in BAT were correlated with higher improvement in RI in 3xTg-AD mice, supporting the idea that higher BAT thermogenesis induced by β3AR stimulation is involved in improved memory performance.

### 3. β3AR stimulation decreases insoluble Aβ42/Aβ40 ratio but has no effect on tau phosphorylation in the hippocampus of 3xTg-AD mice

The 3xTg-AD model allowed us to probe whether the effect of CL-316,243 are related to changes in AD neuropathology. Despite no change in total tau or Aβ burden, a 27% decrease in insoluble Aβ42/Aβ40 ratio was observed in the hippocampus of old 3xTg-AD mice following CL-316,243 injections. This observation is important because Aβ42/Aβ40 ratio in the brain has been consistently associated with higher risk of developing AD, at least in genetic cases. Indeed, the Aβ42/Aβ40 ratio is increased in familial forms of AD and is inversely correlated with the age of onset of the disease (Kumar-Singh et al., 2006; Tanzi, 2012). Decreased Aβ42/Aβ40 ratio suggests a shift in APP cleavage from Aβ42 to Aβ40, which is less prone to aggregation than Aβ42. Importantly, increased Aβ42/Aβ40 ratio precedes amyloid plaques formation in the Tg2576 mouse model of AD (Jacobsen et al., 2006). More recent work using induced pluripotent stem cells (iPSC) corroborates this view, indicating that APP or presenilin 1 (PSEN1) mutations, the more likely to cause familial AD, also leads to higher Aβ42/Aβ40 ratio (Arber et al., 2019). The absence of changes on the phosphorylation status of tau following CL-316,243 administration could be interpreted as surprising. Since tau phosphorylation has been repeatedly shown to follow body temperature modulation (Arendt et al., 2003; Planel et al., 2007; Stieler et al., 2011), we could have expected a protective effect of β3AR stimulation. For example, we recently showed that improved BAT thermogenesis through repeated cold exposure protects old 3xTg-AD mice from cold-induced tau phosphorylation (Tournissac et al., 2019). However, the present study design did not fully explore the hypothesis that β3AR stimulation impacts tau phosphorylation, because the animals were kept at room temperature and not exposed to any frank thermoregulatory challenge (i.e. acute 24-h exposure to cold). Thus, because it has not been directly tested in the present study, it remains possible that pharmacological β3AR stimulation also confers protection against cold-induced tau phosphorylation.

### 4. Conclusion: potential translation to clinic

The present work aimed to determine whether β3AR agonists exert positive effects on AD-relevant endpoints in an animal model of tau and Aβ neuropathology, thereby providing arguments for drug repurposing in AD. This class of drugs is actively being tested in clinical studies for metabolic diseases and CL-316,243 has been previously investigated in humans as well (Weyer et al., 1998). As an example, mirabegron (Myrbetriq^®^) has been approved by the Food and Drug Administration for the treatment of overactive bladder (Chapple et al., 2014b; Cypess et al., 2015), and large randomized controlled trials have confirmed its safety and tolerability profiles (Chapple and Siddiqui, 2017). Thus, the potential translation to clinical use of this class of drugs in AD is high. Nonetheless, it has to be noted that in humans, potential side effects of β3AR agonists include cardiovascular dysfunction induced by non-specific activation of β1 and β2 adrenergic receptors. Indeed, one administration of mirabegron at a dose of 200 mg (fourth times the clinical dose) enhances BAT activity in adults, but also induced cardiac arrhythmia in a few cases (Cypess et al., 2015). While the clinical dose of 50 mg does not seem to be efficient to acutely stimulate BAT (Baskin et al., 2018), a recent study showed that 100 mg of mirabegron enhances thermogenesis without any cardiovascular side effects in adults (Loh et al., 2018). Yet, long-term studies investigating chronic effect of β3AR agonists in BAT thermogenesis in old volunteers are needed.

Altogether, our results in a mouse model of AD demonstrate for the first time that β3AR agonists are potent tools to reverse memory deficits and insoluble Aβ42/Aβ40 ratio in the hippocampus. It is the first study to our knowledge to investigate the potential of this class of drugs in AD neuropathology and behavior.

## AUTHOR CONTRIBUTIONS

M.T. designed the study, performed animal experiments, drug administration, behavior and metabolic evaluations, protein extractions, western blots, immunohistochemistry, HPLC, completed the statistical analysis, interpretation of the data, and wrote the manuscript. T.V. contributed to animal experiments and western blots. N.V. performed western blots and contributed to protein extractions. C.H. and F.P. performed chronobiology analysis on body temperature and reviewed the manuscript. C.T. performed Aβ quantification. K.M. contributed to norepinephrine extraction and measurement. E.P. provided expertise for the tau analysis and reviewed the manuscript. F.C. secured funding, contributed to the experimental design, and wrote the manuscript. F.C. is the guarantor of this work and, as such, had full access to all the data in the study and takes responsibility for the integrity of the data and the accuracy of the data analysis.

## ACKNOWLEDGMENTS

M.T. was funded by a scholarship from the Alzheimer Society of Canada. This study was made possible by funding from the Alzheimer Society of Canada (#<u>15-02</u> and #20-01), the Canadian Institutes of Health Research (<u>MOP 102532</u>), the Quebec Network for Research on Aging and the Canadian Foundation for Innovation (#<u>34480</u>). F.C. and E.P are a Fonds de Recherche du Québec - Santé (FRQ-S) scholars (#<u>253895</u>, #<u>26936</u>, and #<u>252178</u>). The authors are grateful to Patrick Caron and Chantal Guillemette for technical support and to Isabelle Guisle for comments on the manuscript. The authors thank the animal facility technicians Sonia Francoeur, Stéphanie Bernard and France Duclos for their help during the animal experiment.

