## Supplemental Information for "Repurposing beta3-adrenergic receptor agonists for Alzheimer’s disease: Beneficial effects on recognition memory and amyloid pathology in a mouse model"

**SUPPLEMENTARY MATERIAL**

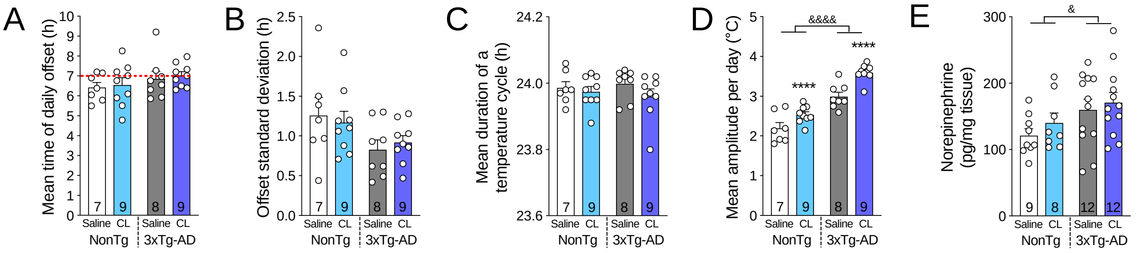


**Figure S1. CL-316,243 administration does not affect circadian rhythm parameters nor BAT norepinephrine content.**

A: individual mean daily offsets (determined as the time of the first six successive bins when temperature was lower than the mean diurnal temperature, thus corresponding to morning temperature drop). B: offset standard deviation. C: mean duration of a total temperature cycle. D: Mean amplitude of body temperature during one day (24-h, from 7 a.m. to 7 p.m.). E: Norepinephrine measured by HPLC in BAT.

Data are represented as mean ± SEM (n/group indicated in bars). Statistics: Two-way ANOVA, effect of treatment: ****p<0.0001, effect of genotype: &p<0.05 &&&&p<0.0001 (A-E).

Abbreviations: 3xTg-AD: triple transgenic mice; NonTg: non-transgenic mice.

| | NonTg | | | | | | 3xTg-AD | | | | | | Two-way ANOVA (p values) | | |
|---|---|---|---|---|---|---|---|---|---|---|---|---|---|---|---|
| | Saline | | | CL-316,243 | | | Saline | | | CL-316,243 | | | Genotype | Treatment | Interaction |
| | Mean | | SD | Mean | | SD | Mean | | SD | Mean | | SD | | | |
| **Synaptic proteins** | | | | | | | | | | | | | | | |
| Drebrin | 2.46 | ± | 0.55 | 2.66 | ± | 0.53 | 2.25 | ± | 0.35 | 2.04 | ± | 0.34 | p=0.0045 | ns | ns |
| PSD95 | 7.17 | ± | 1.59 | 7.03 | ± | 1.27 | 6.86 | ± | 0.61 | 6.41 | ± | 0.66 | ns | ns | ns |
| Septin 3 | 2.19 | ± | 0.93 | 2.13 | ± | 0.76 | 2.21 | ± | 1.02 | 2.23 | ± | 1.14 | ns | ns | ns |
| **Kinases** | | | | | | | | | | | | | | | |
| GSK3β(Ser9) / GSK3β tot | 0.21 | ± | 0.16 | 0.18 | ± | 0.13 | 0.21 | ± | 0.14 | 0.21 | ± | 0.10 | ns | ns | ns |
| AKT (Ser473) / AKT tot | 0.30 | ± | 0.14 | 0.32 | ± | 0.13 | 0.33 | ± | 0.11 | 0.37 | ± | 0.22 | ns | ns | ns |
| **APP production and clearance** | | | | | | | | | | | | | | | |
| BACE-1 | 4.21 | ± | 1.83 | 3.99 | ± | 1.43 | 4.49 | ± | 2.26 | 4.37 | ± | 2.26 | ns | ns | ns |
| APP (6E10) | n/a† | ± | n/a† | n/a† | ± | n/a† | 1.35 | ± | 0.71 | 1.46 | ± | 0.87 | n/a† | ns | ns |
| APP (22C11) | 1.16 | ± | 0.44 | 1.22 | ± | 0.41 | 2.81 | ± | 1.46 | 2.91 | ± | 1.60 | p<0.0001 | ns | ns |
| APP C-terminal fragment | 2.27 | ± | 1.01 | 2.34 | ± | 0.91 | 4.04 | ± | 2.16 | 4.17 | ± | 2.31 | p<0.0001 | ns | ns |
| sAPPα | n/a† | ± | n/a† | n/a† | ± | n/a† | 1.84 | ± | 0.88 | 1.87 | ± | 0.90 | n/a† | ns | ns |
| X11α | 1.98 | ± | 0.47 | 2.07 | ± | 0.42 | 1.32 | ± | 0.28 | 1.28 | ± | 0.33 | p<0.0001 | ns | ns |
| IDE | 2.45 | ± | 0.48 | 2.62 | ± | 0.35 | 2.31 | ± | 0.58 | 2.19 | ± | 0.71 | ns | ns | ns |
| LRP1 | 1.36 | ± | 0.33 | 1.38 | ± | 0.38 | 1.40 | ± | 0.42 | 1.36 | ± | 0.46 | ns | ns | ns |
| RAGE | 1.90 | ± | 0.87 | 1.81 | ± | 0.58 | 2.29 | ± | 1.16 | 2.39 | ± | 1.25 | ns | ns | ns |
| **Others** | | | | | | | | | | | | | | | |
| Bax / Bcl-2 | 11.51 | ± | 7.44 | 10.05 | ± | 5.41 | 10.61 | ± | 3.32 | 12.18 | ± | 5.61 | ns | ns | ns |
| GFAP | 3.10 | ± | 1.26 | 2.81 | ± | 0.95 | 3.53 | ± | 1.73 | 3.89 | ± | 1.86 | ns | ns | ns |
| NeuN | 1.58 | ± | 0.41 | 1.63 | ± | 0.32 | 1.69 | ± | 0.35 | 1.69 | ± | 0.38 | ns | ns | ns |
| GLUT1 50 kDa | 0.64 | ± | 0.20 | 0.59 | ± | 0.38 | 0.36 | ± | 0.20 | 0.36 | ± | 0.11 | p=0.0014 | ns | ns |
| GLUT1 45 kDa | 2.76 | ± | 0.71 | 2.80 | ± | 0.72 | 2.26 | ± | 0.34 | 2.20 | ± | 0.38 | p=0.0022 | ns | ns |

†: non applicable, human-specific antibody

**Table S1. Other AD markers not affected by CL-316,243 treatment.**

Relative optical density of proteins normalized on actin measured in detergent-soluble fraction of hippocampus homogenates by Western Blot. Data are represented as mean ± SD, n= 9-12 per group. Abbreviations: AKT: protein kinase A; APP: amyloid precursor protein; BACE-1: betasecretase 1; GFAP: glial fibrillary acidic protein; GLUT1: glucose transporter 1; GSK3β, glycogen synthase kinase 3β; IDE: insulin degrading enzyme; LRP1: low density lipoprotein receptor related protein 1; PSD95: post-synaptic density 95; RAGE: receptor of advanced glycation end products; sAPPα: soluble α-APP.

| Antibody | Clone | Specificity | Host | Source |
|---|---|---|---|---|
| | | ***Primary antibodies*** | | |
| **Actin** | monoclonal | β-actin | Mouse | Applied Biological Materials (Richmond, BC, Canada) |
| **AKT** | polyclonal | AKT a.a. 345-480 | Rabbit | Santa Cruz Biotechnology Inc. (Santa Cruz, CA, USA) |
| **AKT (phospho Ser473)** | polyclonal | AKT, phosphorylated at Ser-473 | Rabbit | EMD Millipore (Billerica, MA, USA) |
| **Aβ (MOAB-2)** | 6C3 | Amyloid-β 40 and 42, unaggregated, oligomeric and firbillar | Mouse | EMD Millipore (Billerica, MA, USA) |
| **APP/Aβ (6E10)** | 6E10 | APP a.a. 1-16 | Mouse | Covance, Inc. (Princeton, NJ, USA) |
| **APP (22C11)** | monoclonal | All three forms of APP: immature, sAPP and mature | Mouse | EMD Millipore (Billerica, MA, USA) |
| **APP CTF** | polyclonal | a.a. 751-770 of APP | Rabbit | EMD Millipore (Billerica, MA, USA) |
| **sAPPα** | 2B3 | Synthetic peptide of the C-terminal part of Human sAPPα | Mouse | IBL (Fujioka, Japan) |
| **β3AR** | polyclonal | mouse beta 3 adrenergic receptor, aa 350 to C-teminus | Rabbit | Abcam (Cambridge, MA, USA) |
| **BACE-1** | monoclonal | synthetic peptide (ab108394) | Mouse | EMD Millipore (Billerica, MA, USA) |
| **Bax** | polyclonal | total bax protein | Rabbit | Cell Signaling Technology (Danvers, MA, USA) |
| **Bcl-2** | polyclonal | total bcl-2 alpha protein | Rabbit | Cell Signaling Technology (Danvers, MA, USA) |
| **Drebrin** | Mx823 | c-term peptide (a.a.632-649) | Mouse | Progen Biotechnik GmbH (Heidelberg, Germany) |
| **eEF2** | polyclonal | total eEF2 protein independant of phosphorylation | Rabbit | Cell Signaling Technology (Danvers, MA, USA) |
| **GFAP** | GA-5 | GFAP | Mouse | Sigma-Aldrich (St.Louis, MO, USA) |
| **GLUT1** | monoclonal | synthetic peptide, c-terminus (ab40084) | Mouse | Abcam (Cambridge, MA, USA) |
| **GSK3β** | monoclonal | Rat GSK-3*β* aa. 1-160 | Mouse | BD Biosciences (Mississauga, ON, Canada) |
| **GSK3β (phospho Ser9)** | polyclonal | GSK3β, phosphorylated at Ser-9 | Rabbit | Cell Signaling Technology (Danvers, MA, USA) |
| **IDE** | polyclonal | a.a. 93-273 | Rabbit | Abcam (Cambridge, MA, USA) |
| **LRP1** | EPR3724 | synthetic peptide | Rabbit | Abcam (Cambridge, MA, USA) |
| **Mitochondrial oxidative phosphorylation system** | monoclonal | Total OXPHOS complexes: CI, CII, CIII, CIV and CV subunits | Rabbit | Abcam (Cambridge, MA, USA) |
| **NeuN** | monoclonal | a.a. 1-100 | Rabbit | Abcam (Cambridge, MA, USA) |
| **PSD95** | monoclonal | aa 77-299 | Mouse | NeuroMab (Davis, CA, USA) |
| **RAGE** | monoclonal | Gly23 - Leu342 | Rat | R&D system (Minneapolis, MN, USA) |
| **Septin3** | polyclonal | synthetic peptide | Rabbit | Novus biologicals (Oakville, ON, Canada) |
| **Tau (total) TauC** | polyclonal | c-terminal region of tau protein | Rabbit | Dako (Burlington, ON, Canada) |
| **Tau (total)** | Tau13 | Total human tau | Mouse | Covance, Inc. (Princeton, NJ, USA) |
| **Tau (phospho)** | Tau1-PC1C6 | Tau, phosphorylated at Ser195, Ser198, Ser199 and Ser202 | Mouse | EMD Millipore (Billerica, MA, USA) |
| **Tau (phospho)** | AT180 | Tau, phosphorylated at Thr-231 | Mouse | Pierce Endogen Inc. (Rockford, IL, USA) |
| **Tau (phospho)** | AT270 | Tau, phosphorylated at Thr-181 | Mouse | Pierce Endogen Inc. (Rockford, IL, USA) |
| **Tau (phospho)** | CP13 | Tau, phosphorylated at Ser-202 | Mouse | Generous gift from Peter Davies |
| **Tau (phospho)** | PHF1 | Tau, phosphorylated at Ser-396 and Ser-404 | Mouse | Generous gift from Peter Davies |
| **UCP1** | monoclonal | a.a. 145-159 | Rabbit | Abcam (Cambridge, MA, USA) |
| **X11α** | polyclonal | X11, a.a. 1-220 | Rabbit | Santa Cruz Biotechnology Inc. (Santa Cruz, CA, USA) |
| | | ***Secondary antibodies*** | | |
| **Goat anti-mouse** | | Peroxidase-conjugated AffiniPure Goat anti-mouse IgG (H+L) | Goat | Jackson ImmunoResearch (West Grove, PA, USA) |
| **Goat anti-rabbit** | | Peroxidase-conjugated AffiniPure Goat anti-rabbit IgG (H+L) | Goat | Jackson ImmunoResearch (West Grove, PA, USA) |

**Table S2. Antibodies used in this study.**
